# Tau accumulates in Crohn’s disease gut

**DOI:** 10.1101/2020.03.16.979534

**Authors:** Alice Prigent, Guillaume Chapelet, Adrien de Guilhem de Lataillade, Thibauld Oullier, Emilie Durieu, Arnaud Bourreille, Emilie Duchalais, Kévin Hardonnière, Michel Neunlist, Wendy Noble, Saadia Kerdine-Römer, Pascal Derkinderen, Malvyne Rolli-Derkinderen

## Abstract

A sizeable body of evidence has recently emerged to suggest that gastrointestinal inflammation might be involved in the development of Parkinson’s disease. There is now strong epidemiological and genetical evidence linking Parkinson’s disease to inflammatory bowel diseases and we recently demonstrated that the neuronal protein alpha-synuclein, which is critically involved in Parkinson’s disease pathophysiology, is upregulated in inflamed segments of Crohn’s colon. The microtubule associated protein tau is another neuronal protein critically involved in neurodegenerative disorders but, in contrast to alpha-synuclein, no data are available about its expression and phosphorylation patterns in inflammatory bowel diseases. Here, we examined the expression levels of tau isoforms, their phosphorylation profile and truncation in colon biopsy specimens from 16 Crohn’s disease and 6 ulcerative colitis patients and compared them to samples from 16 controls. Additional experiments were performed in full thickness segments of colon of 5 Crohn’s disease and 5 control subjects, in primary cultures of rat enteric neurons and in Nrf2 knockout mice. Our results show the upregulation of two main human tau isoforms in the enteric nervous system in Crohn’s disease but not in ulcerative colitis. This upregulation was not transcriptionally regulated but instead likely resulted from a decrease in protein clearance via an Nrf2 pathway. Our findings, which provide the first detailed characterization of tau in Crohn’s disease, suggest that the key proteins involved in neurodegenerative disorders such as alpha-synuclein and tau, might also play a role in Crohn’s disease.

## Introduction

Crohn’s disease (CD) is a type of inflammatory bowel disease, which is characterized by chronic remitting and relapsing inflammation of the entire gastrointestinal (GI) tract. Although the precise etiology of CD is unknown, it is generally accepted that it develops through an exaggerated immune response to luminal antigens in genetically susceptible individuals leading to intestinal epithelium inflammation and damage (1). The pathological process in CD is however not limited to the epithelial lining but extends to all components of the GI wall including the enteric nervous system (ENS) whose structure and neurochemical phenotype are altered during active CD (2). The major structural abnormalities thus far described in CD include architectural alterations of the plexuses, nerve hypertrophy and hyperplasia along with alterations of neuronal cell bodies (3, 4). Regarding neurochemical phenotype, the two most comprehensive studies found an increased number of vasoactive intestinal polypeptide-immunoreactive nerve cell bodies in both the myenteric and submucosal plexuses (5, 6).

The ENS structure and neurochemistry resemble that of the CNS, therefore pathogenic mechanisms that give rise to CNS disorders might also lead to ENS dysfunction, and *vice versa* (7). Over the past two decades, progress in molecular genetics and neuropathology enabled a better understanding of the pathogenesis of neurodegenerative disorders. The observation that abnormal protein accumulation is characteristic of particular disease sets has led to a neuropathological classification according to the composition of the abnormal protein aggregates. The most common neurodegenerative disorders are now classified based on the abnormal accumulation of alpha-synuclein (8) or tau (9) and hence referred to as synucleinopathies (with Parkinson’s disease being the prototypical type) or tauopathies (with Alzheimer’s disease being the most common), respectively (10, 11). It should be noted however that the pathological distinction among these disorders is sometimes not so clear as the co-occurrence of alpha-synuclein and tau deposits has been reported in PD (12). Several independent studies have shown that enteric neurons, like their CNS counterparts, also express alpha-synuclein (13–15) and tau (16, 17) and we recently demonstrated that alpha-synuclein is upregulated in inflamed segments of Crohn’s colon (18). These latter findings, together with the results of recent epidemiological and genetical studies (reviewed in (19)), support the existence of a close relationship between Parkinson’s and Crohn’s disease. In contrast to alpha-synuclein, no data are available about the expression and phosphorylation patterns of enteric tau in CD. Here, we examined the expression levels of tau isoforms, their phosphorylation profile and truncation in colon biopsy specimens from CD patients and compared them to samples from controls. Additional pharmacological experiments were performed in primary cultures of rat enteric neurons. Our results show the upregulation of two main human tau isoforms in the ENS in CD, which results from a decrease in protein clearance via an Nrf2 pathway. Our findings, which provide the first detailed characterization of tau in CD, suggest that the key proteins involved in neurodegenerative disorders such as alpha-synuclein and tau, might also play a role in CD, and provide mechanistic insights into the events underlying tau and alpha-synuclein accumulation in CD gut.

## Material and methods

### Participants and human tissues

A total of 38 subjects participated in this study, 16 CD and 6 ulcerative colitis (UC) patients as well as 16 healthy controls who had a normal colonoscopy for colorectal cancer screening (supplemental table 1; All 6 UC patients, 10/16 CD patients and 10/16 controls from the current study had their biopsies previously analyzed in (18) for the expression levels of alpha-synuclein). The study protocol was approved by the local Committee on Ethics and Human Research (*Comité de Protection des Personnes Ouest VI*). Written informed consent was obtained from each patient and from each normal volunteer before the endoscopic procedure. All procedures were performed according to the guidelines of the French Ethics Committee for Research on Humans and registered under the no. DC-2008-402. CD and UC patients had 4 biopsies (2 in an inflamed and 2 in a non-inflamed area) taken during the course of a colonoscopy and stored in RA1 buffer (Macherey Nagel, Hoerdt, France) until further analysis. For controls, 6 biopsies were taken and also stored in RA1 buffer. The inflammatory status of CD and UC biopsies was confirmed by qPCR for TNF-α and IL1-β (supplemental figure 1).

Samples from 5 additional CD patients and controls were used for immunohistochemistry and sarkosyl extraction experiments. FFPE full thickness segments of colon containing the myenteric plexus were obtained from 3 CD patients and controls subjects who underwent colonic resection for disease management and non-obstructive colorectal carcinoma, respectively. Frozen full thickness segments of colon from two additional CD patients and controls were used for sarkosyl extraction experiments.

Samples of frozen temporal cortex from two post-mortem human brains, one with Alzheimer’s disease (AD) and one devoid of neurodegeneration were obtained from the Neuropathology Department of Angers (Dr Franck Letournel) to serve as a control for the following experiments.

### Primary culture of rat ENS

Primary cultures of rat ENS were generated and cultured as previously described (Coquenlorge et al., 2014). Cells were treated with 10 µM sulforaphane or MG132 for 12h.

### Knockout mice

Nrf2−/− mice were provided by the RIKEN BRC according to an MTA to Prof S. Kerdine-Römer (20). Mice were housed in a pathogen-free facility and handled in accordance with the principles and procedures outlined in Council Directive 86/609/EEC. Genotyping was performed by PCR using genomic DNA that was isolated from tail snips as described (20).

### Western Blot

Two biopsies per patient stored in RA1 buffer were pooled and lysed using the “Precellys 24” tissue homogenizer (Bertin technologies, St Quentin-en-Yvelines, France) followed by sonication with “vibracell 75 186” device (Sonics, Newton CT, USA). Protein and RNA extractions were performed with NucleoSpin Triprep Kit (Macherey-Nagel, Hoerdt, France) according to the manufacturer’s instructions. The total proteins were precipitated and prepared for Polyacrylamide Gel Electrophoresis using protein precipitator and resuspension buffer PSB/TCEP (Protein solving buffer and (tris(2-carboxyethyl)phosphine) TCEP reducing agent) from the NucleoSpin Triprep Kit (Macherey-Nagel, Hoerdt, France). The elution fraction containing RNA was further used for PCR analysis (see below). Cells (primary culture of rat ENS) and tissues from mice (hippocampus, ileum, proximal and distal colon) were lysed in RIPA lysis buffer (Merck Millipore, Fontenay sous Bois, France) containing phosphatase inhibitor cocktail II (Roche, Neuilly sur Seine, France) and protease inhibitors cocktail (Roche). Total protein was quantified using a Nanodrop 2000 spectrophotometer (ThermoFisher Scientific, Cillebon sur Yvette, France). Equal amounts of tissue or cell lysates were separated and analyzed by Western blot as described previously using Invitrogen NuPage Novex 10% Bis Tris MiniGels™ (18). Tau protein ladder (Sigma) was used in some experiments as a control. The primary antibodies used are listed in Table 1. Mouse monoclonal anti-PGP 9.5 (1:2,000; Ultraclone limited, Isle of Wight, UK) antibody was used as a pan-neuronal marker and membranes were reprobed with mouse monoclonal anti-β-actin antibody (1:10,000; Sigma, Saint-Quentin Fallavier, France) to confirm equal protein loading. For quantification, the relevant immunoreactive bands were quantified with laser-scanning densitometry and analyzed with NIH Image J software. To allow comparison between different films, the density of the bands was expressed as a percentage of the average of controls.

**Table 1.**
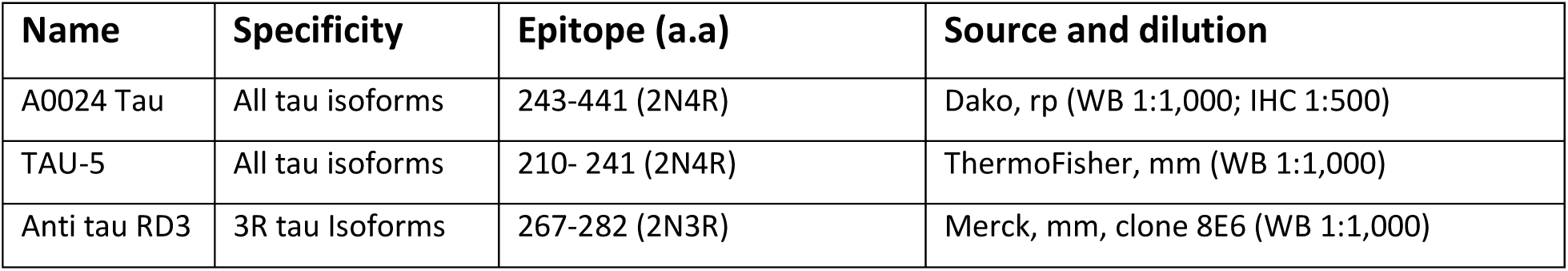
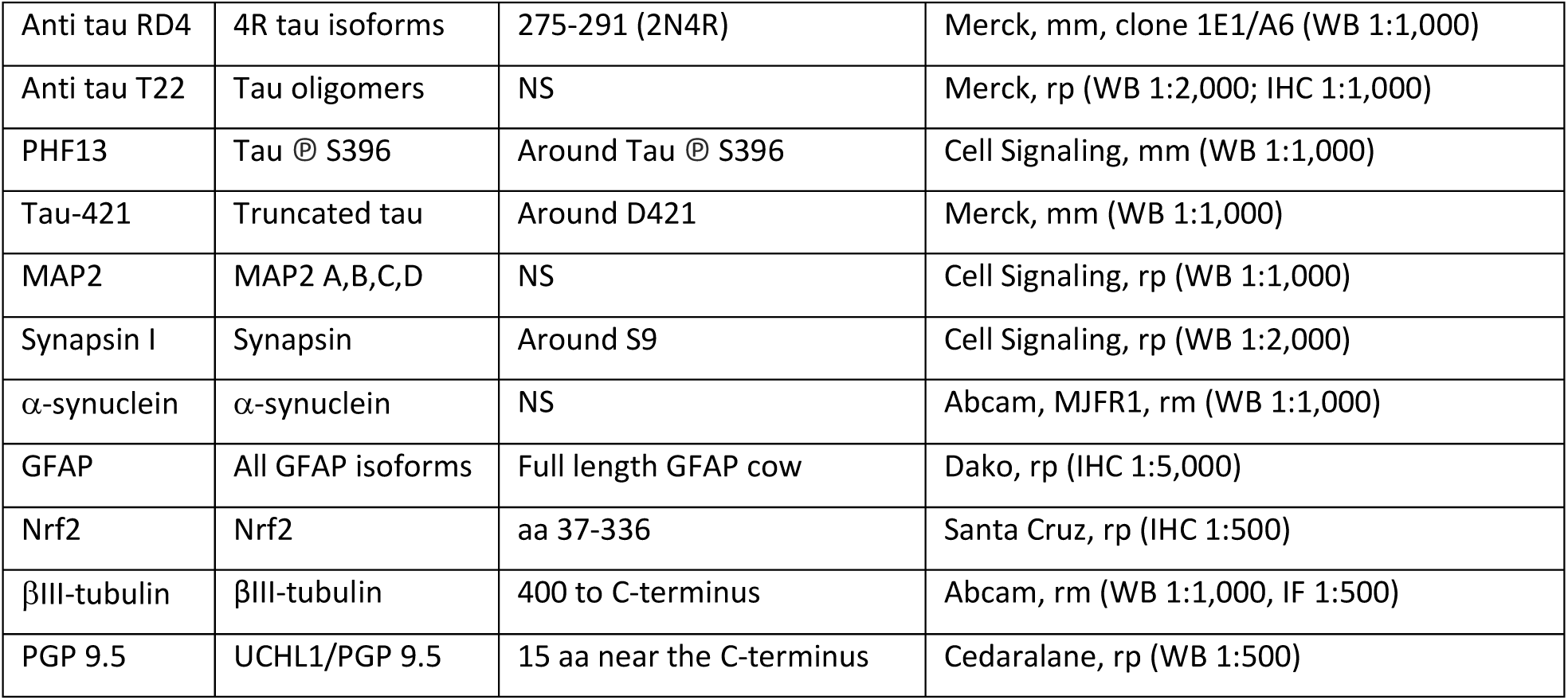
The name, specificity, epitope, source and dilution of the antibodies used in this study are shown. Abbreviations are amino-acids (aa); immunohistochemistry (IHC); mouse monoclonal (mm); not specified (NS); rm (rabbit monoclonal); rabbit polyclonal (rp); western blot (WB).

### Immunohistochemistry

Fixed human tissues from 3 CD patients and 3 controls were included in paraffin using an embedding station (MICROM MICROTECH STP120) and sections (5µm) were performed using a microtome (MICROM HM355S). The slides were deparaffinized with 2 xylene baths (for 5 min each) and incubated in 4 ethanol baths (100%, 95%, 70%, 50%, respectively for 3 min each). After a rinse in distilled water, slides were washed in PBS and antigen retrieval was performed using a Sodium Citrate solution (2.94g Sodium Citrate Tribase; 1 L distilled water; 500 µL Tween 20; pH 6) at 95°C for 20 min. Slides were incubated in NH4Cl (100mM) for 15 min before incubation in PBS containing 0.5% (v/v) Triton X-100 for 1 hour then in 10 % (v/v) horse serum in PBS with 0.5% (v/v) Triton X-100. Primary antibodies (Table 1) were incubated overnight at 4°C, then with secondary antibody at room temperature for 2 hours. Images were acquired with a fluorescent microscope AxioZoom.V16 (Zeiss, Marly le Roi, France) associated with Zen 2012 software (Zeiss). Analyses were performed with the ImageJ software (NIH, Bethesda, MD). Fluorescence immunohistochemical signals of n = 18 and 17 myenteric ganglia were recorded from each specimen of the control and CD group respectively. Each myenteric ganglion was marked and the mean grey value (integrated density/ μm^2^ ganglionic tissue) was determined.

### Sarkosyl extraction and ultracentrifugation

For sarkosyl extraction of tau, tissues were homogenized in 50 mM Tris buffered saline (TBS, pH 7.4) containing 2 mM EGTA, 1 mM Na_3_VO_4_, 10 mM sodium fluoride, 1 mM phenylmethylsulfonyl fluoride and 10 % (w/v) sucrose (all from Sigma). Samples were centrifuged at 12,000 g for 20 min (4 °C) to remove cells debris and 10% sarkosyl (v/v) was added to the resultant supernatant to give a final concentration of 1% (v/v). Samples were mixed for 30 min at ambient temperature 4°C, then centrifuged at 100,000 g for 1 h at 21°C. The supernatant (sarkosyl-soluble tau) was collected and the pellet (sarkosyl-insoluble tau) was washed twice with 1% (v/v) sarkosyl. Proteins in the samples were resolved by Polyacrylamide Gel Electrophoresis and processed for immunoblotting as described in the Western Blot section. Samples of frozen temporal cortex from one post-mortem human brain with Alzheimer’s disease (AD) and one control subject were obtained from the Neuropathology Department of Angers (Dr Franck Letournel) to serve as a control.

### RT-PCR

RNA extraction was performed with NucleoSpin Triprep Kit (Macherey-Nagel, Hoerdt, France) in two-pooled biopsies (see Western Blot section) according to the manufacturer’s instructions. One μg purified mRNA was denatured and processed for reverse transcription using Superscript III reverse transcriptase (Thermo Fisher Scientific, Saint-Herblain, France). PCR amplifications were performed using the Absolute Blue SYBR green fluorescein kit (Roche Molecular Biochemicals, Meylan, France, Cat# AB4166B) and run on a StepOnePlus system (Life Technologies, Cat# 4376600). The following primers were used:

- MAPT, forward: 5’ AGAGTCCAGTCGAAGATTGGGTC 3’; reverse: 5’ GGGTTTCAATCTTTTTATTTCCTCC 3’
- SNCA, forward: 5’ CCAAAGAGCAAGTGACAAATGTTG 3’; reverse: 5’ AGCCAGTGGCTGCTGCAAT 3’
- TNF-α, forward: 5’ CCCGAGTGACAAGCCTGTAG 3’; reverse: 5’ TGAGGTACAGGCCCTCTGAT 3’
- IL-1β, forward: 5’ GAGCAACAAGTGGTGTTCTCC 3’; reverse: 5’ TTGGGATCTACACTCTCCAGC 3’
- RPS6, forward: 5’ AAGCACCCAAGATTCAGCGT 3’; reverse: 5’ TAGCCTCCTTCATTCTCTTGGC 3’

### Statistics

All data shown are mean ± SEM. Statistical analyses were performed using GraphPad software version 8.00 (San Diego California, USA). Between-group comparisons were performed using an independent sample t test (Mann-Whitney test), Kruskal-Wallis or Chi-squared test. Differences were deemed statistically significant when p<0.05.

## Results

The main clinical features of the study population are shown in supplementary table 1. CD patients were significantly younger than controls (supplemental table 1).

### Tau expression is increased in Crohn’s disease but not in ulcerative colitis

We first analyzed the expression levels of tau in colonic biopsies of CD and controls by Western blot. A significant 8- and 4-fold increase in tau protein expression was observed in the inflamed area of CD patients when compared to controls, after normalization to PGP 9.5 and β-actin, respectively (Figure 1a and supplemental figure 2). By contrast, no significant differences were observed when the non-inflamed area was compared to controls (Figure 1a and supplemental figure 2) and when biopsies of UC patients, either in the inflamed or non-inflamed area were compared to controls (supplemental figure 2). We have recently shown that 1N3R and 0N4R are the two main tau isoforms expressed in the adult human ENS and that these two isoforms are phosphorylated under physiological conditions (16). Using 3R and 4R isoform specific antibodies, we found that both 1N3R and 0N4R tau isoforms were upregulated in the inflamed area in CD (Figure 1b). We next examined the phosphorylation state of tau with a phospho-specific antibody that detect tau phosphorylated at Ser396. After normalizing to total tau, the amount of phosphorylated tau was not different between CD samples and controls (Figure 1c). Truncated tau is present in the pathological deposits observed in CNS tauopathies, including PSP and Alzheimer’s disease (21). In order to determine whether tau is C-terminally truncated in the enteric neurons in CD, colonic biopsies were analyzed using a monoclonal antibody that specifically recognizes tau cleaved at Asp421 by caspase-3. After normalizing to total tau, the amount of truncated tau was dramatically decreased in CD inflamed area when compared to either non-inflamed area or controls (Figure 1d). Additional immunohistochemistry experiments were performed to examine the localization and the expression levels of tau in the myenteric plexus. Colonic myenteric plexus showed intense tau immunoreactivity in both neuronal cell bodies and processes, which nearly completely overlapped beta-tubulin immunostaining (Figure 2a). Quantification of the immunofluorescence showed increased tau immunoreactivity in CD myenteric plexus as compared to controls (Figure 2b).

**Figure 1.**
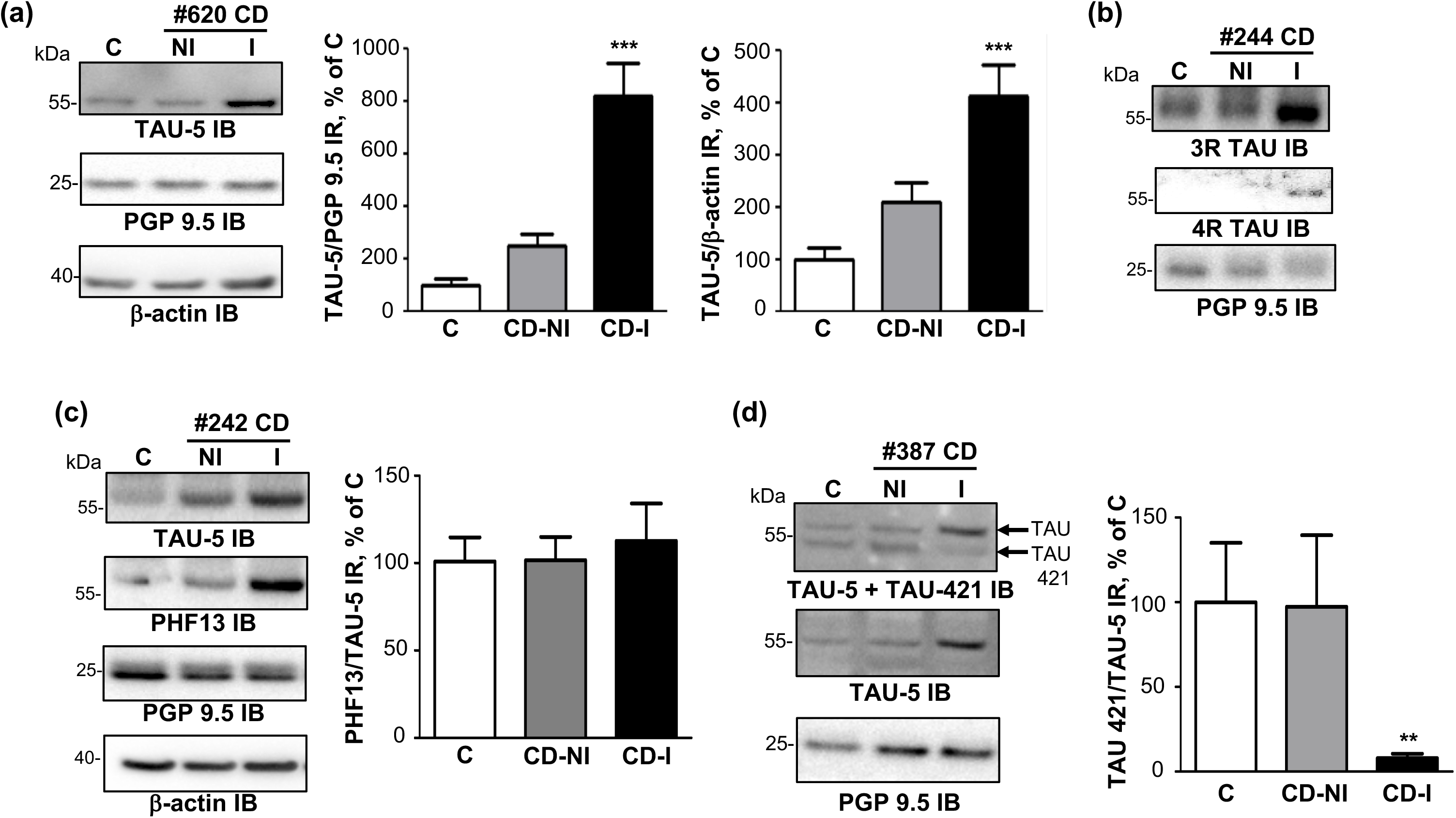
Expression levels, phosphorylation state and truncation of tau protein in colonic biopsies from patients with Crohn’s disease. **(a)** Colonic biopsy lysates from 16 Crohn’s disease (CD) patients and 16 controls (C) (see supplemental table 1 for subject details) were immunoblotted with total tau (TAU-5 IB), PGP 9.5 (PGP 9.5 IB) and β-actin (β-actin IB) antibodies (raw immunoblots are provided in supplemental figure 2). For CD patients, biopsies taken in non-inflammatory (NI, NI-CD) and inflammatory areas (I, I-CD) were analyzed separately. Tau-immunoreactive bands were measured, normalized to the optical densities of PGP 9.5 and β-actin and expressed as percentage of controls. Data correspond to mean ± SEM of 16 samples for CD patients control subjects (C), *** p<0.001 **(b)** Representative immunoblots showing the expression of 3R and 4R tau using specific isoforms antibodies (8 CD and 8 controls were analyzed) **(c)** Colonic biopsies lysates from 16 CD patients and 16 controls (C) were immunoblotted with total tau (TAU-5 IB), tau phosphorylated at serine 396 (PHF13 IB), PGP 9.5 (PGP 9.5 IB) and β-actin (β-actin IB) antibodies. Phosphorylated tau immunoreactive bands were measured, normalized to the optical densities of total tau and expressed as percentage of controls. Data correspond to mean ± SEM of 16 samples for CD patients and control subjects **(d)** Colonic biopsies lysates from 16 CD patients and 16 controls (C) were immunoblotted with total tau (TAU-5 IB), truncated tau at Asp 421 (TAU-421) and PGP 9.5 (PGP 9.5 IB) antibodies. In the upper panel, the membrane was probed with both antibodies. TAU-421 immunoreactive bands were measured, normalized to the optical densities of total tau and expressed as percentage of controls. Data correspond to mean ± SEM of 16 samples for CD patients and control subjects, ** p<0.01.

**Figure 2.**
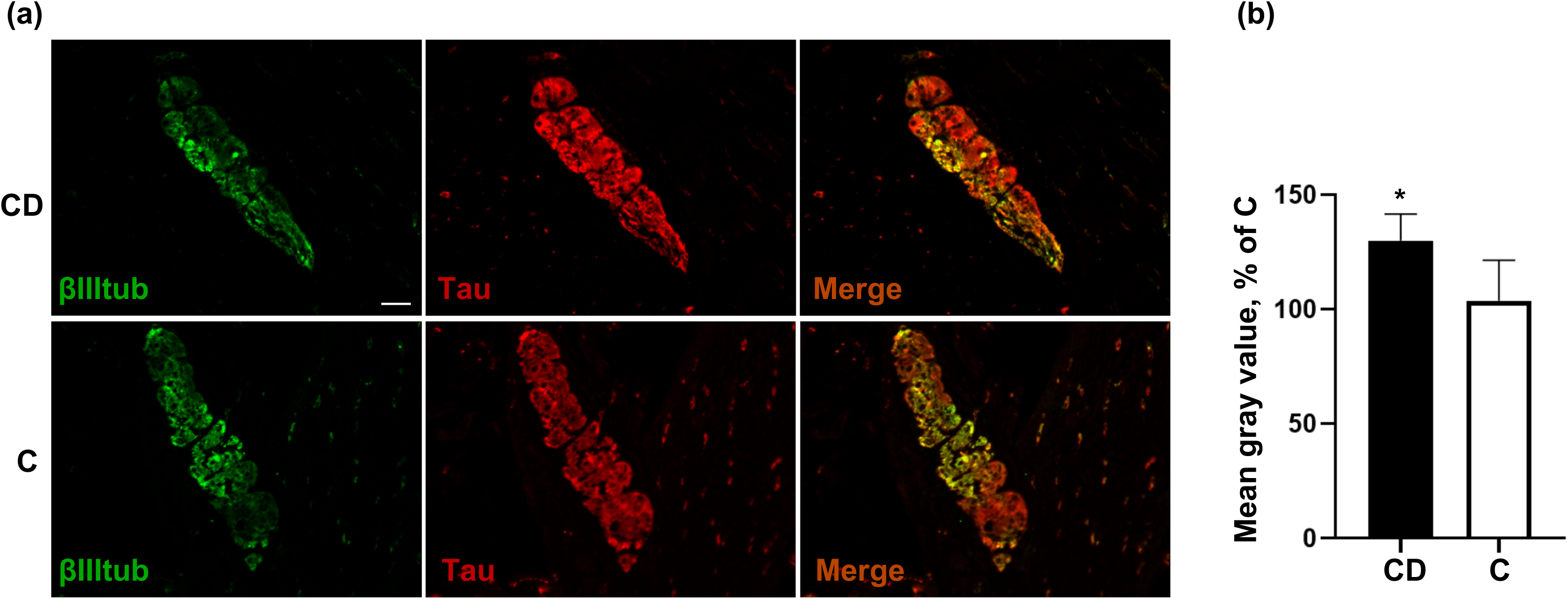
Localization and expression levels of tau in the myenteric plexus in Crohn’s disease and controls. **(a)** Total tau antibody A0024 (Tau) was used to detect tau in the myenteric ganglia in CD and control subjects (C); beta III-tubulin (βIII-tub) antibody was used to specifically label neurons. Scale bar is 50 μm **(b)** Quantitative analysis of tau immunofluorescence in the myenteric plexus of CD and controls (C). Data are shown as corrected mean gray value normalized to controls and presented as mean ± SEM, n=18 and 17 myenteric ganglia for control and CD, respectively, **p < 0.01.

### The upregulation of tau in Crohn’s disease is not accompanied with more insoluble or oligomeric forms of the protein

Tau cleaved by caspase-3 at Asp421 is more prone to aggregation and acts as a nidus for aggregation of full-length tau (22, 23). To examine whether tau upregulation is associated with protein aggregation, we used T22, a recently developed tau oligomer-specific antibody (24). As expected, this antibody detected slower migrating tau bands in AD brain when compared to control brain (Figure 3a). However, no difference in the banding pattern was observed when this antibody was used to compare biopsy lysates from CD and controls (Figure 3b). In CNS tauopathies, the biochemical characterization of pathological tau involves well-standardized protocols based on the use of sarkosyl (25)(Figure 3c). Using this method along with the T22 antibody, we showed that the levels of sarkosyl-insoluble tau was increased in AD when compared to control brain (Figure 3d). In contrast, no sarkosyl-insoluble tau was detected in the GI tract samples from either CD patients or control subjects (Figure 3d).

**Figure 3.**
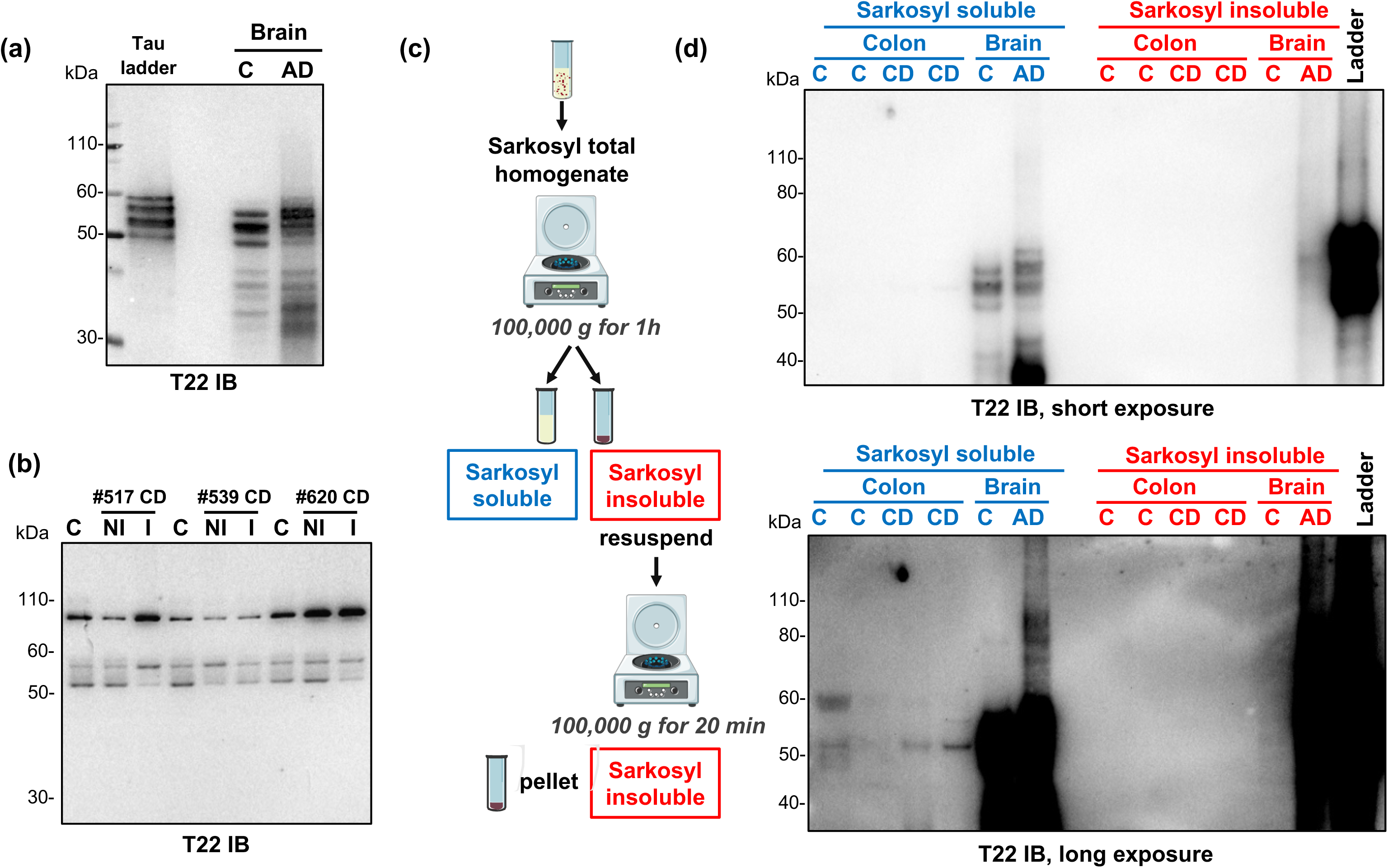
Analysis of sarkosyl-insoluble tau and oligomeric forms of tau in colonic biopsies from patients with Crohn’s disease. **(a)** Lysates from control (C) and Alzheimer’s disease (AD) brains served as controls to ensure that the immunoreactive pattern observed with T22 was different in AD brain when compared to control brain. **(b)** Colonic biopsy lysates from 3 CD patients (#517, 539 and 620) and 3 controls (C) were subjected to immunoblot analysis using Tau T22 antibody **(c and d)** Full thickness segments of colon from 2 CD and 2 controls along with samples of control (C) and Alzheimer’s disease (AD) brains were separated into sarkosyl-soluble and insoluble fractions as described in the methods and in the diagram. Tau was analyzed by immunoblotting using Tau T22.

### Tau upregulation in Crohn’s disease is specific as not observed for MAP2c, βIII-tubulin and synapsin I

To determine whether the upregulation of tau is specific to this microtubule associated protein, we analyzed the expression levels of its sequence homologue MAP2c along with two of its interacting proteins, βIII-tubulin and synapsin I (26, 27). Unlike tau, the amounts of MAP2c and βIII-Tubulin were not different between CD and controls (Figure 4). By contrast, synapsin I expression was significantly decreased in samples of CD colon when compared to controls (Figure 4).

**Figure 4.**
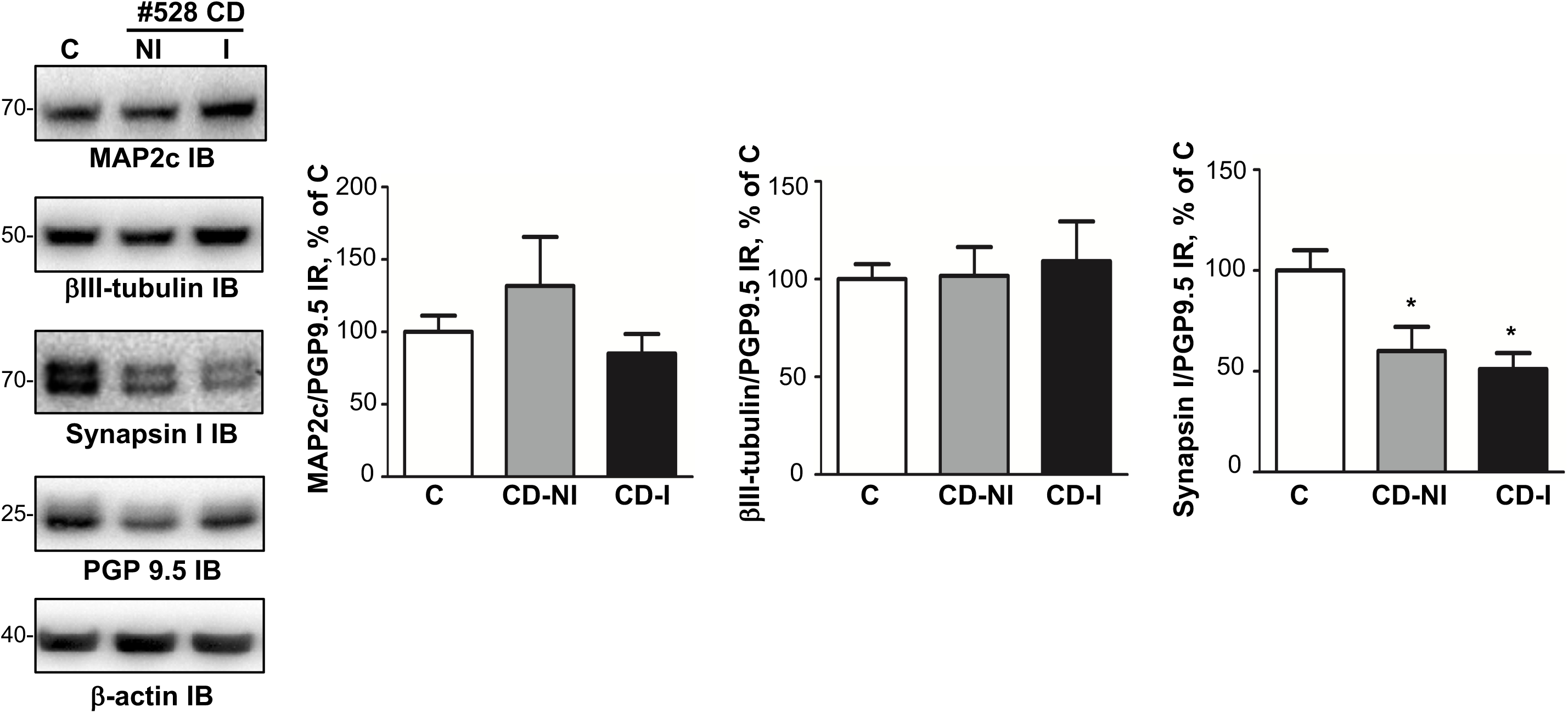
Expression levels of MAP2c, synapsin I and β-III tubulin in colonic biopsies from patients with Crohn’s disease. Colonic biopsy lysates from 6 CD patients (#242, 243, 244, 248, 528 and 539) and 6 controls (C) were immunoblotted with MAP2c (MAP2c IB), synapsin I (Synapsin I IB) and β-III tubulin (β-actin IB) antibodies. MAP2c, synapsin I and β-III tubulin immunoreactive bands were measured, normalized to the optical densities of PGP9.5 and expressed as percentage control. Data correspond to mean ± SEM of 6 samples for CD patients and control subjects (*p<0.05)

### The increase in tau expression likely results from a deregulated Nrf2/NDP52 pathway

We next sought to determine the mechanisms by which tau is upregulated in CD. Using RT-PCR, we found a decrease in tau mRNA in inflamed-CD samples as compared to controls (Figure 5a). In addition, we did not observe any difference between CD and controls when samples were analyzed by Western Blot for the amounts of TRIM 28, a transcriptional regulator that modulates both alpha-synuclein and tau levels (28) (Figure 5b). These results demonstrate that the higher amount of tau observed in CD does not result from an increase in tau transcription and suggest that other mechanisms such as modulation in tau proteostasis might be involved. A sizeable body of evidence have shown that tau could be degraded through the proteasome/ubiquitin pathway or turned over by the autophagy-lysosome pathway (29). We first investigated the possible role of the proteasome/ubiquitin pathway by analyzing ubiquitin immunoreactivity in whole biopsy lysates by Western blot. The banding pattern was markedly different between CD and controls, with some stronger immunoreactive bands around 25, 35 and 50-60 kDa being observed in inflamed CD samples (supplemental figure 3). To further investigate the possible contribution of the proteasome to the degradation of tau in enteric neurons, we examined the levels of tau in primary culture of rat ENS in the presence of the proteasome inhibitor MG132. In line with our previous results, primary culture of rat ENS express four tau isoforms (0N3R, 1N3R/0N4R and 2N3R), which migrate as a triplet of 50, 53 and 58 kDa bands on SDS PAGE (16). Treatment with MG132 decreased the amount of all these tau isoforms (supplemental figure 3) thereby suggesting that proteasome inhibition is not involved in tau upregulation in enteric neurons.

**Figure 5.**
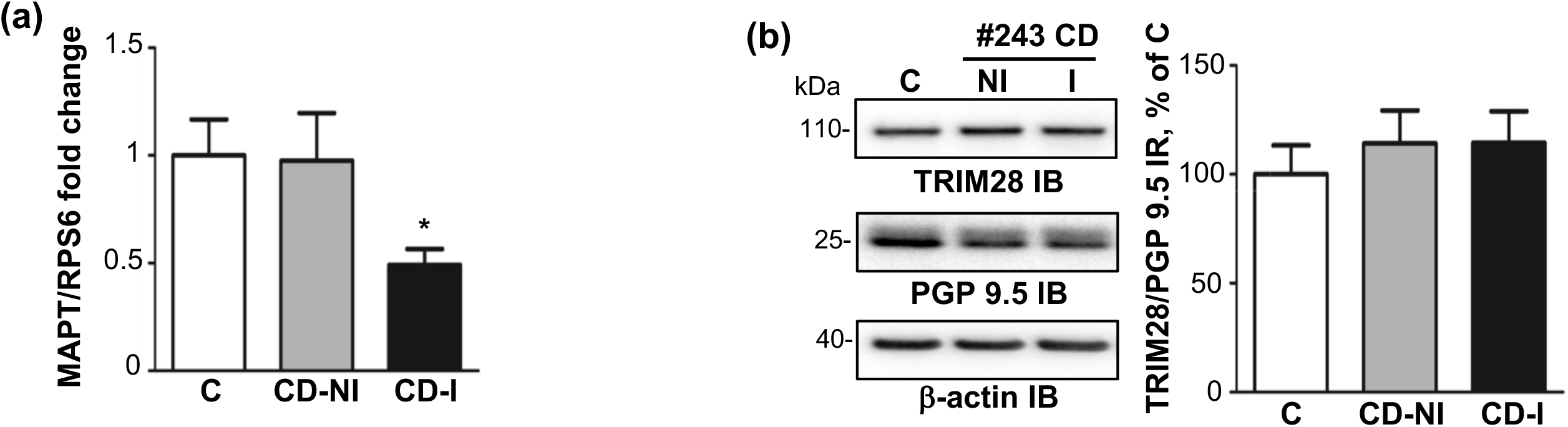
Transcriptional regulation of tau in Crohn’s disease. **(a)** Quantitative PCR analysis of tau mRNA (MAPT) in colonic biopsy from all 16 CD patients (CD-NI and CD-I) and all 16 controls. Values represent mean ± SEM (n=16 samples per condition; *p<0.05) **(b)** Colonic biopsy lysates from all 16 CD patients and all 16 controls were immunoblotted with antibodies against TRIM28 (TRIM28 IB), PGP 9.5 (PGP 9.5 IB) and β-actin (β-actin IB). TRIM28 immunoreactive bands were measured, normalized to the optical densities of PGP 9.5 and expressed as a percentage of the average of controls.

A recent study showed that the nuclear factor erythroid 2-related factor 2 (Nrf2) decreased the levels of tau by inducing the autophagy adaptor protein NDP52 (also known as CALCOCO2) in CNS neurons (30). When we analyzed the expression of Nrf2 in the ENS, we found that its amount was significantly decreased in CD inflamed area (Figure 6). A parallel decrease of NDP52 was also observed (Figure 6). To further investigate the possible role of Nrf2, we used the Nrf2 activator sulforaphane in primary culture of ENS. Nrf2 is primarily expressed by enteric glial cells and to a lesser extent by enteric neurons (Figure 7a) and treatment with sulforaphane induced a significant decrease in the expression levels of all tau isoforms (Figure 7b), without changing its phosphorylation status (Figure 7b). We next examined the levels of total and phosphorylated tau in the GI tissues of Nrf2-knockout mice. As shown in Figure 7c, a 2-fold increase in tau protein expression was observed in the distal colon of 21-week-old Nrf2 knockout mice in comparison with wild-type mice. No changes were observed in the distal colon of 11-week-old mice (Figure 7c). Tau phosphorylation levels were similar between Nrf2 knockout mice in comparison with wild-type mice, regardless of age (Figure 7c).

**Figure 6.**
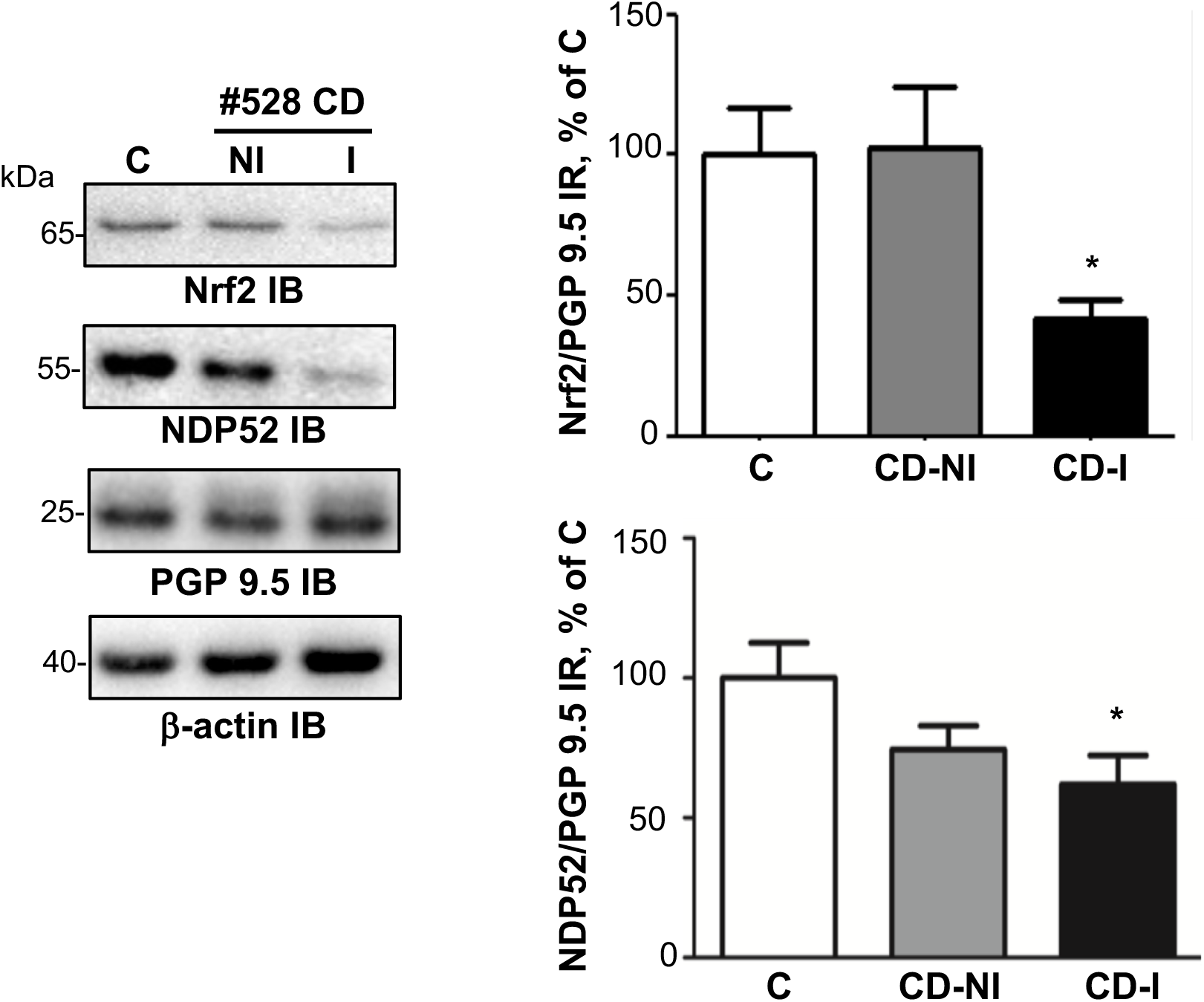
Nrf2 and NDP52 expression in Crohn’s disease. **(a)** Expression levels of Nrf2 in CD. Colonic biopsy lysates from 8 CD patients (#242, 243, 244, 248, 517, 528, 539, 620) and 8 controls (C) were immunoblotted with Nrf2 (Nrf2 IB), NDP52 (NDP52 IB), PGP 9.5 (PGP 9.5 IB) and β-actin (β-actin IB) antibodies. Nrf2 and NDP52immunoreactive bands were measured, normalized to the optical densities of PGP9.5 and expressed as percentage control. Data correspond to mean ± SEM of 8 samples for CD patients and control subjects (*p<0.05)

**Figure 7.**
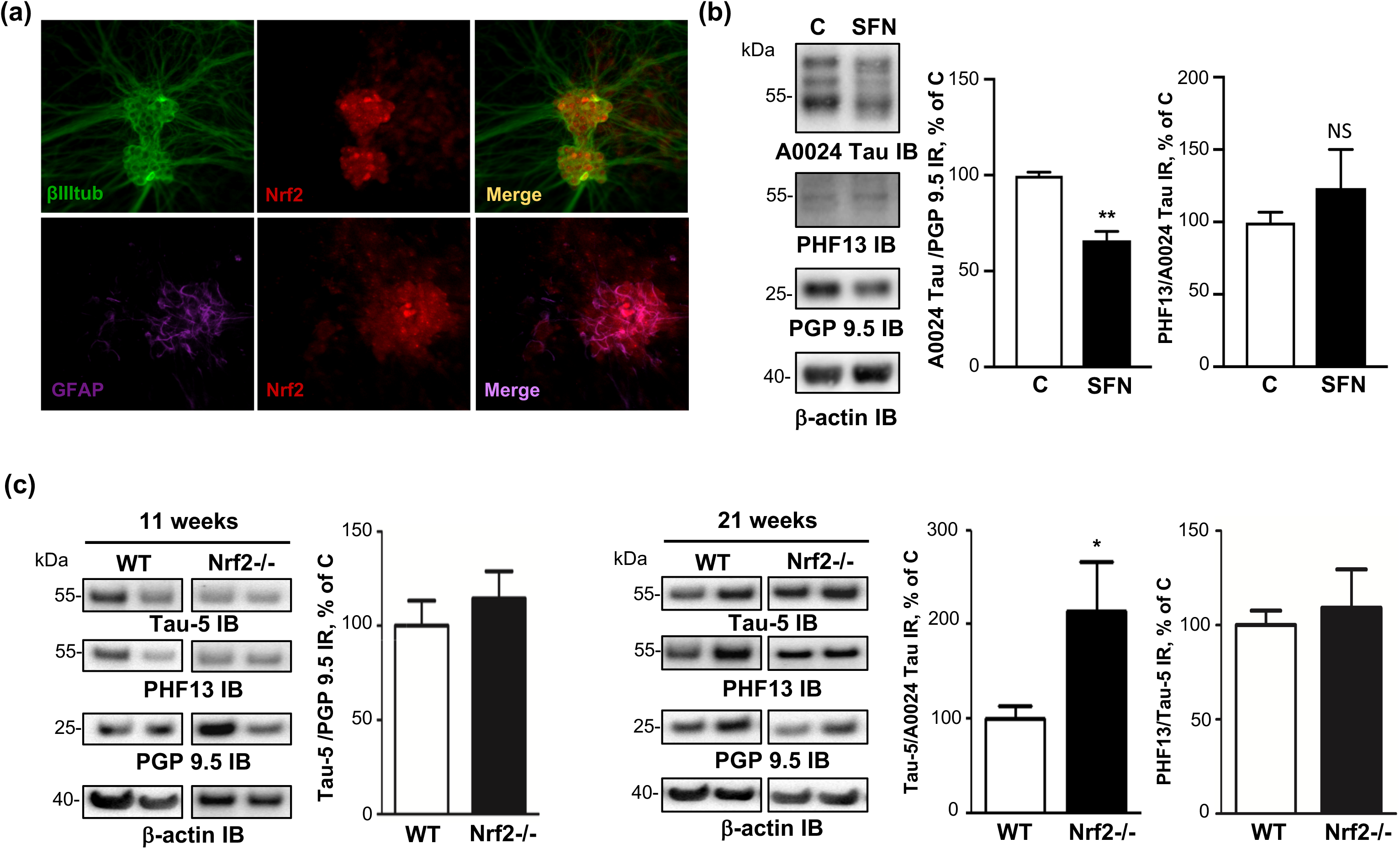
Role of Nrf2 in the regulation of enteric tau expression. **(a)** Distribution and localization of Nrf2 in primary ENS culture. Primary culture of rat ENS at 14 days *in vitro* were immunolabeled with anti-Nrf2, anti-βIII tubulin antibodies. Scale bar is 100 µm for the upper panel and 50 µm for the lower panel **(b)** Effects of sulforaphane on the expression levels of tau in primary culture of rat ENS. After 14 days in culture, primary culture of rat ENS were treated with vehicle (PBS, control) or 10 µM sulforaphane. Cell lysates were immunoblotted with total tau (A0024 Tau IB), tau phosphorylated at serine 396 (PHF13 IB), PGP 9.5 (PGP 9.5 IB) and β-actin (β-actin IB) antibodies. Total tau and phosphorylated tau immunoreactive bands were measured, normalized to the optical densities of PGP9.5 and total tau, respectively and expressed as percentage of controls. Values represent mean ± SEM (n=8; treated versus control; **p<0.01) **(c)** Distal colon lysates obtained from Nrf2-/- (11-week-old, two male and one female; 21-week-old, three male) or wild-type (WT, 11 months old, three male; 21-week-old, one female and two male) mice were immunoblotted with total tau (TAU-5 IB), tau phosphorylated at serine 396 (PHF13 IB), PGP 9.5 (PGP 9.5 IB) and β-actin (β-actin IB) antibodies. Total tau and phosphorylated tau immunoreactive bands were measured, normalized to the optical densities of PGP9.5 and total tau, respectively and expressed as percentage of controls.

### Alpha-synuclein and tau are similarly regulated in CD

We previously used the colonic samples from 10/16 CD patients and 10/10 controls to show that alpha-synuclein was upregulated in CD inflamed area (18). We extend these previous findings by showing that alpha-synuclein mRNA expression is down-regulated in CD (Figure 8a) and that it correlated with the amount of tau mRNA (Figure 8b). Such a positive correlation was also observed at the protein level (Figure 8c). Finally, we showed that, as with tau, treatment of primary culture of ENS with sulforaphane induced reduction of alpha-synuclein and that the expression of alpha-synuclein was higher in the distal colon of 21-week-old Nrf2 knockout mice when compared to wild-type mice (Figure 7d)

**Figure 8.**
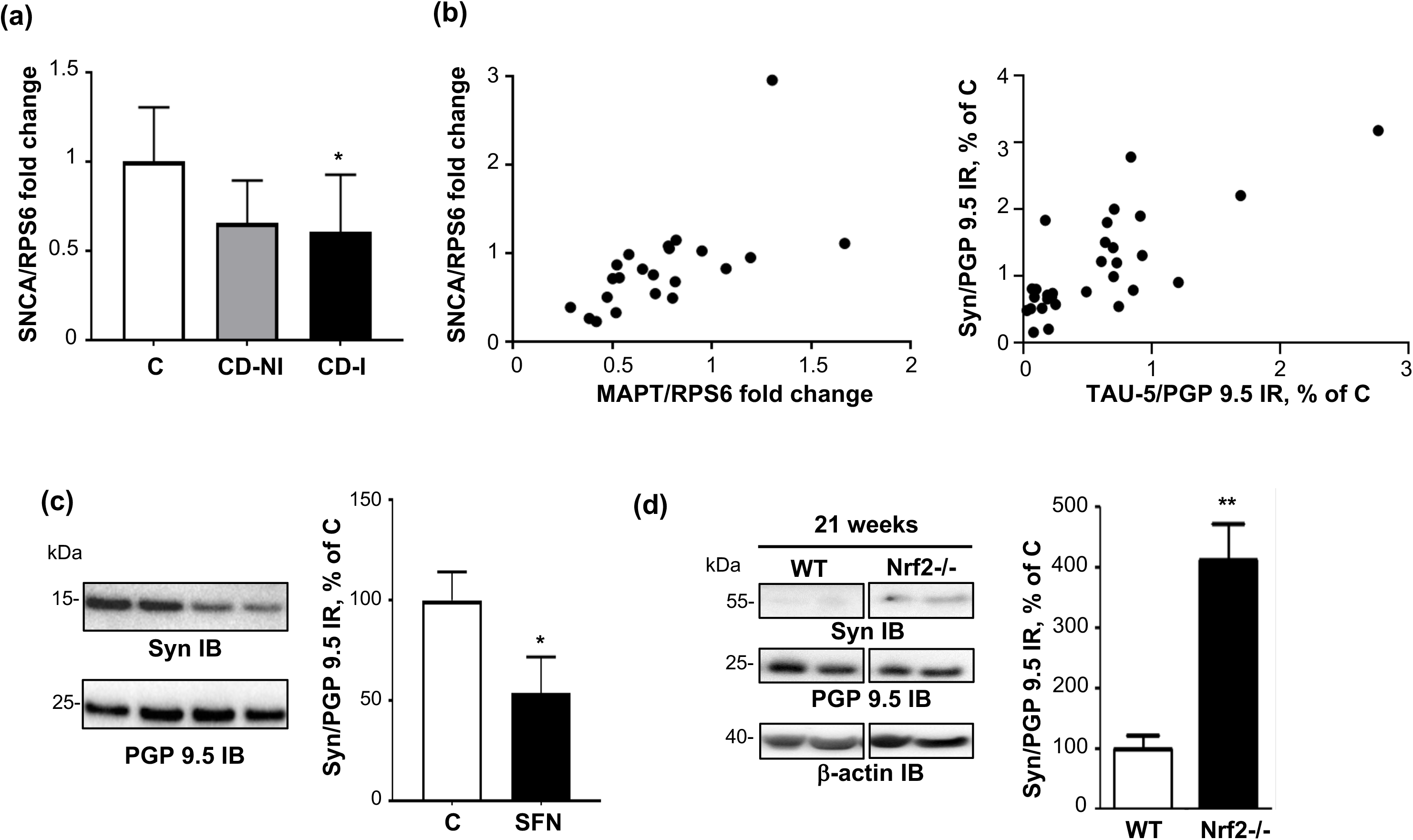
Expression level of enteric alpha-synuclein and its correlation with tau. **(a)** Quantitative PCR analysis of alpha-synuclein mRNA (SNCA) in colonic biopsy from 7 CD patients (CD-NI and CD-I, #248, 381, 409, 461, 539, 620) and 7 controls (#191, 194, 330, 335, 345, 377 and 477) **(b)** Correlation between SNCA and MAPT mRNA expression levels in colonic biopsy from 7 CD patients (CD-NI and CD-I, #248, 381, 409, 461, 539, 620) and 7 controls (#191, 194, 330, 335, 345, 377 and 477) (r=0.73; p<0.0001) **(c)** Correlation between alpha-synuclein and tau protein expression levels in colonic biopsy from 10 CD patients (CD-NI and CD-I, #381, 387, 409, 436, 458, 461, 516, 517, 533 and 620) and 10 controls (#330, 335, 344, 345, 359, 377, 428, 430, 450 and 456) (r=0.67; p<0.0001) **(d)** Effects of sulforaphane on the expression levels of alpha-synuclein in primary culture of rat ENS. After 14 days in culture, primary culture of rat ENS were treated with vehicle (PBS, control) or 10 µM sulforaphane. Cell lysates were immunoblotted with alpha-synuclein (A-Syn IB), PGP 9.5 (PGP 9.5 IB) and β-actin (β-actin IB) antibodies. Alpha-synuclein immunoreactive bands were measured, normalized to the optical densities of PGP9.5 and expressed as percentage of controls. Values represent mean ± SEM (n=6; treated versus control; *p<0.05) **(e)** Distal colon lysates obtained from Nrf2-/- (21-week-old, three male) or wild-type (WT; 21-week-old, one female and two male) mice were immunoblotted with alpha-synuclein (Syn IB), PGP 9.5 (PGP 9.5 IB) and β-actin (β-actin IB) antibodies. Alpha-synuclein immunoreactive bands were measured, normalized to the optical densities of PGP9.5 and expressed as percentage of controls.

## Discussion

We have recently shown that the isoform profile of tau differed between the ENS and the CNS, with 1N3R and 0N4R being the two main tau isoforms expressed in adult human enteric neurons (16). In addition, we did not observe any changes in tau expression, phosphorylation or truncation of colonic samples from patients suffering from the prototypical tauopathy progressive supranuclear palsy (PSP). These results, along with the findings from a recent comprehensive autopsy survey (17) strongly suggest that unlike PD, the pathological process in tauopathies such as Alzheimer’s disease and PSP is limited to the CNS and does not involve the ENS. We therefore set out to examine the expression levels and phosphorylation state of tau in CD, an inflammatory bowel disease that has been consistently associated with ENS abnormalities (4). We found that the expression levels of enteric tau isoforms are dramatically increased in the colon of CD patients and that this upregulation is likely mediated through the Nrf2/NDP52. By contrast, no changes in the expression levels of tau was observed in UC.

The ENS may be involved in inflammatory bowel disease as a result of tissue injury, or via the effects of soluble mediators of the inflammatory process including cytokines, arachidonic acid metabolites and oxygen-derived free radicals. Although changes in the morphology of the ENS have been described in both CD and UC, structural changes are more pronounced in CD than in UC (reviewed in (4, 31)). In this study, synapsin I, a neuronal phosphoprotein associated with the cytoplasmic surface of synaptic vesicles, was significantly decreased in samples of CD colon. These findings are in line with previous observations that showed severe and extensive submucosal axonal necrosis in the ileum and colon in CD but not in UC (32). This axonal necrosis may also explain the observed decrease in tau mRNAs, which are transported along the axon to be translated *in situ* in the axon terminals (33).

Numerous proteases, such as aminopeptidases, calpains and caspases have been shown to proteolyze tau, however, most of these enzymes do not appear to be principally responsible for tau clearance. It is therefore unlikely that the sole decrease in tau truncation at Asp 421 in CD is sufficient to induce overexpression. The bulk of clearance of both physiological and pathological forms of tau is instead mediated by the proteasomal and autophagic degradative systems (29, 34). While several reports suggested that tau could be degraded through the proteasome, others did not confirm this observation and even observed increased tau clearance in the presence of proteasome inhibitors in primary cortical neurons (30, 35). Our findings obtained in primary culture of ENS, which showed a decreased in tau in the presence of proteasome inhibitor MG132 are therefore in line with this latter observation. Although we could not exclude completely a role for other pathways, our data strongly indicate that the Nrf2/NDP52 autophagic pathway is involved in the accumulation of tau in CD. As the Nrf2 signaling system plays a key role in the maintenance of cellular homeostasis under stress, inflammatory, carcinogenic, and pro-apoptotic conditions, it seems quite logical to observe a decreased expression of Nrf2 in CD (36). It should be however mentioned that, as far as we know, our study is the first to show that Nrf2 is downregulated in inflammatory bowel disease. Our observation reinforce the findings obtained in rodents, in which activating and/or upregulating Nrf2 alleviated DSS-induced colitis (37, 38), thereby suggesting than reinforcing Nrf2 could be a potential therapeutic option for inflammatory bowel disease patients.

Hyperphosphorylation, truncation and aggregation of tau are characteristic features of the pathological deposits observed in CNS tauopathies (34). Using specific antibodies and sequential extraction/ultracentrifugation, we did not find evidence of tau pathological forms in the inflamed zone in CD (we even observed a decrease in the amount of truncated tau). These findings are in keeping with the ones we obtained with alpha-synuclein, which is overexpressed but not aggregated in the ENS in CD (18, 39). One obvious limitation of our work is that the neuronal density in the ENS is rather low and that most of the analyses for the current study were essentially restricted to the analysis of the submucosal plexus. We can therefore not rule out that the absence of overt pathological changes in tau in colonic samples from our CD patients may be due to this limited regional analysis and perhaps different findings would have been obtained had we examined larger GI samples or used more sensitive approach such as protein misfolding cyclic amplification (40).

Our current and previous findings demonstrate that both tau and alpha-synuclein, which are critically involved in neurodegenerative disorders are upregulated in CD further reinforcing the possible link between inflammatory bowel disease and neurodegenerative disorders (19). Some recent observations not only showed that PD and inflammatory bowel disease are epidemiologically (41) but also genetically linked. The Leucine-rich repeat kinase 2 (LRRK2) gene, which has emerged as the gene most commonly associated with both familial and sporadic PD, has been subsequently identified by as a major susceptibility gene for CD (42). Quite interestingly, molecular and neuropathological links between tau and LRRK2 have also been demonstrated: (i) LRRK2 binds to and phosphorylates tau (43, 44) (ii) tau is a prominent pathology in LRRK2 Parkinson’s disease as well as in mice that overexpress mutated forms of LRRK2 (45, 46) (iii) LRRK2 promotes tau accumulation and aggregation and enhances the neuronal transmission of tau in the mouse brain (43, 47). It remains however to be determined why only a subset of patients with inflammatory bowel disease and gastrointestinal inflammation will eventually develop a neurodegenerative disorder. A recent opinion paper has proposed that PD pathogenesis can be divided into three temporal phases: ‘triggers’, which set off the disease process in the brain and/or peripheral tissues, ‘facilitators’ that help triggers access the nervous system or spread the pathology within the CNS and ‘aggravators’, which may for instance increase alpha-synuclein spreading (48). In such a scenario, can be suggested that gastrointestinal inflammation may act as a facilitator.

## List of abbreviations

AD: Alzheimer’s disease
CD: Crohn’s disease
EGTA: ethylene glycol-bis(β-aminoethyl ether)-N,N,N′,N′-tetraacetic acid
ENS: Enteric nervous system
GFAP: Glial fibrillary acidic protein
GI: Gastrointestinal
NDP52: Nuclear dot protein 52 kDa
Nrf2: Nuclear factor erythroid 2-related factor
PD: Parkinson’s disease
TBS: Tris buffered saline
UC: Ulcerative colitis

## Acknowledgements and sources of support

This work was supported by *centre d’entraide et de coordination des associations de parkinsoniens* (CECAP) and *Parkinsonien de Vendée* (FFGP) to AP, AGL and PD.

## Authors contributions

P. Derkinderen, M. Rolli-Derkinderen, Michel Neunlist and S. Kerdine-Römer designed research; A. Prigent, G. Chapelet, A. de Lataillade, T. Oullier, E. Durieu and K. Hardonnière performed research; A. Bourreille and E. Duchalais supervised biobanking; P. Derkinderen, M. Rolli-Derkinderen and W. Noble wrote the paper.

