## Supplemental Table 1 for "Tau accumulates in Crohn’s disease gut"

| Patient # | Diagnosis | Age | Sex | Treatment |
|---|---|---|---|---|
| **242** | CD | 23 | F | IMS |
| **243** | CD | 52 | F | cortico + 5-ASA |
| **244** | CD | 31 | M | IMS |
| **248** | CD | 45 | F | antiTNF |
| **381** | CD | 34 | M | IMS |
| **387** | CD | 43 | F | antiTNF |
| **409** | CD | 34 | M | antiTNF |
| **436** | CD | 20 | M | antiTNF |
| **458** | CD | 38 | F | None since 2 years |
| **461** | CD | 35 | M | antiTNF |
| **516** | CD | 39 | M | antiTNF - IMS |
| **517** | CD | 22 | M | antiTNF |
| **528** | CD | 21 | F | 5-ASA |
| **533** | CD | 24 | M | antiTNF - IMS |
| **539** | CD | 39 | M | antiTNF+ IMS |
| **620** | CD | 38 | F | None since 2 years |
| **283** | UC | 31 | F | 5-ASA |
| **429** | UC | 45 | M | IMS |
| **438** | UC | 36 | F | 5-ASA |
| **494** | UC | 37 | F | antiTNF |
| **504** | UC | 49 | M | antiTNF - 5-ASA |
| **586** | UC | 38 | M | antiTNF - 5-ASA |
| **190** | Control | 51 | M | - |
| **191** | Control | 64 | M | - |
| **192** | Control | 70 | F | - |
| **193** | Control | 72 | F | - |
| **194** | Control | 63 | F | - |
| **195** | Control | 61 | F | - |
| **330** | Control | 55 | M | - |
| **335** | Control | 73 | F | - |
| **344** | Control | 41 | F | - |
| **345** | Control | 48 | F | - |
| **377** | Control | 48 | F | - |
| **430** | Control | 41 | F | - |
| **450** | Control | 49 | M | - |
| **456** | Control | 47 | F | - |
| **460** | Control | 31 | F | - |
| **477** | Control | 18 | F | - |

**Supplemental table 1.** Demographic data for patients and controls. CD: Crohn’s disease; UC: ulcerative colitis.

| | Total (n= 38) | Control (n = 16) | CD (n=16) | UC (n=6) | p value * (control Vs CD patients) | p value ** (3 groups) | p value *** (control vs UC patients) | p value **** (UC vs CD patients |
|---|---|---|---|---|---|---|---|---|
| **Age, mean +/- SD** | 42.3 +/- 14.5 | 52.0 +/- 15.1 | 33.6 +/- 9.5 | 33.9 | **< 0.001** | **0.001** | 0.064 | 0.294 |
| **Female, n (%)** | 22 (57.9) | 12 (75.0) | 7 (43.8) | 3 (50.0) | 0.149 | 0.184 | 0.334 | 0.793 |

* Between-group comparisons (**Control vs CD patients**) were performed using an independent sample t test or Chi-squared test ($\mathcal{X}2$), as appropriate. bold values are significant

** Between-group comparisons (**3 groups**) were performed using an independent sample Kuskal-Wallis test or Chi-squared test ($\mathcal{X}2$), as appropriate. bold values are significant

*** Between-group comparisons (**Control vs UC patients**) were performed using an independent sample t test or Chi-squared test ($\mathcal{X}2$), as appropriate. bold values are significant

**** Between-group comparisons (**CD vs UC patients**) were performed using an independent sample t test or Chi-squared test ($\mathcal{X}2$), as appropriate. bold values are significant
