## Supplemental Figures for "Tau accumulates in Crohn’s disease gut"

**Supplementary file 1**

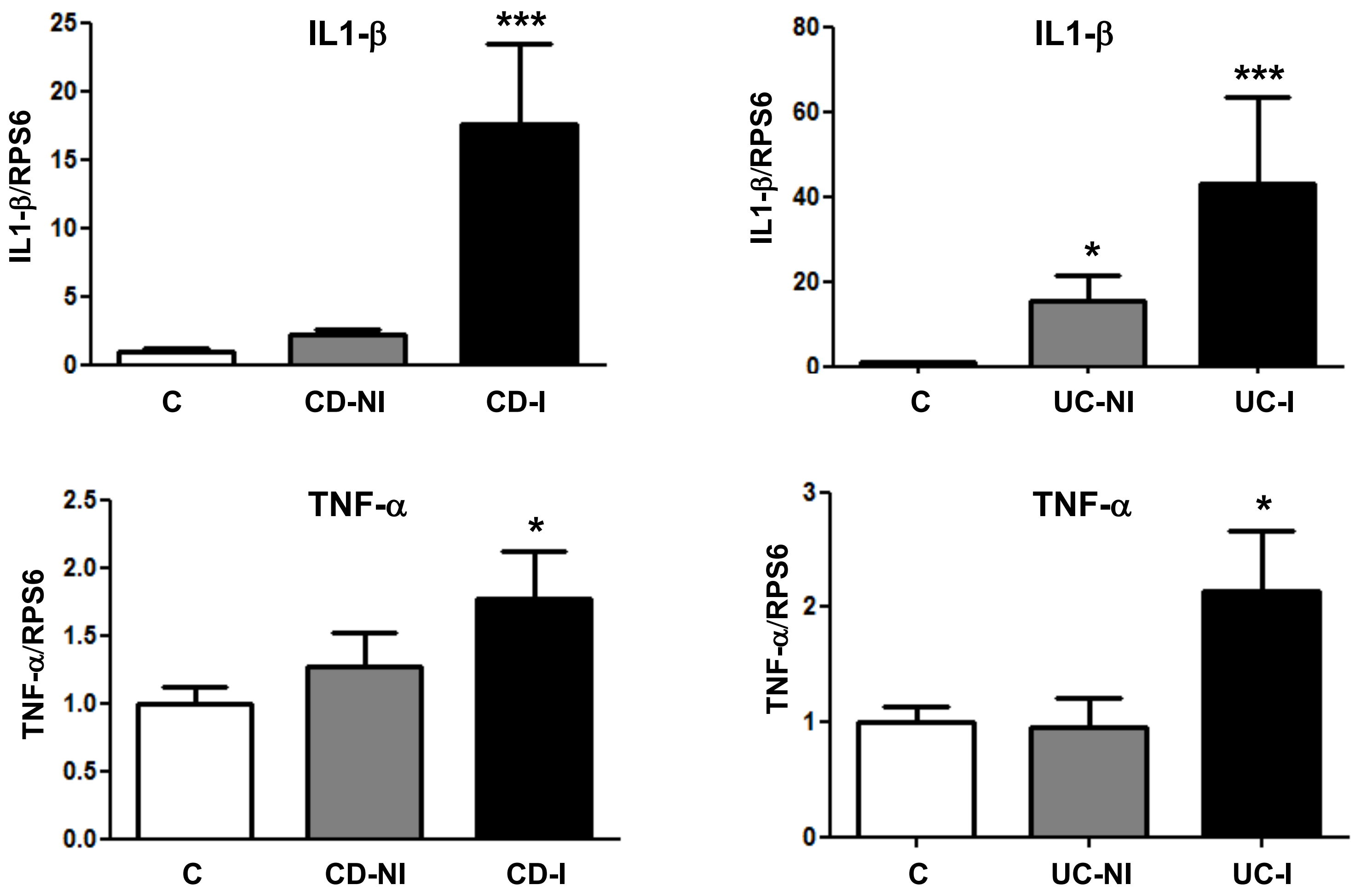


Quantitative PCR analysis of IL1-β and TNF-α mRNA in colonic biopsy from all 16 CD patients (CD-NI and CD-I), all 6 UC patients (UC-NI and UC-I) and all 16 controls. Values represent mean ± SEM (*p<0.05 ; ***p<0.001)

### Supplementary file 2

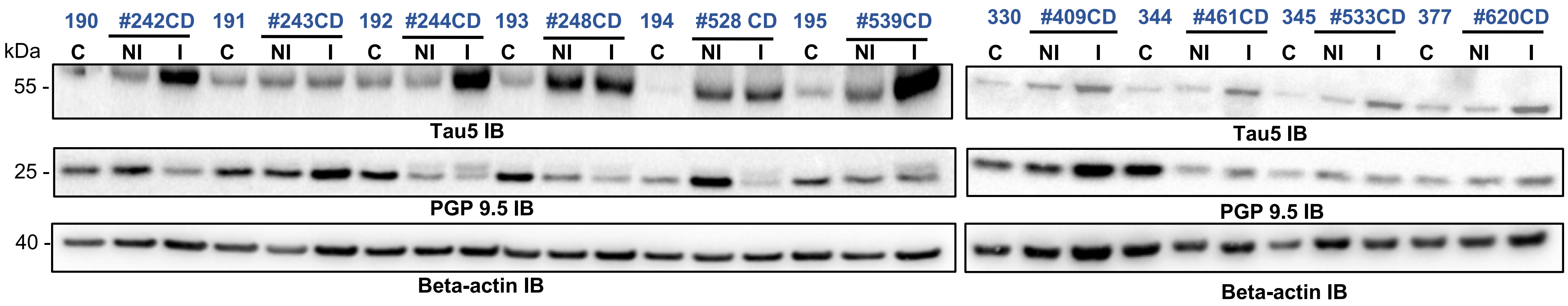


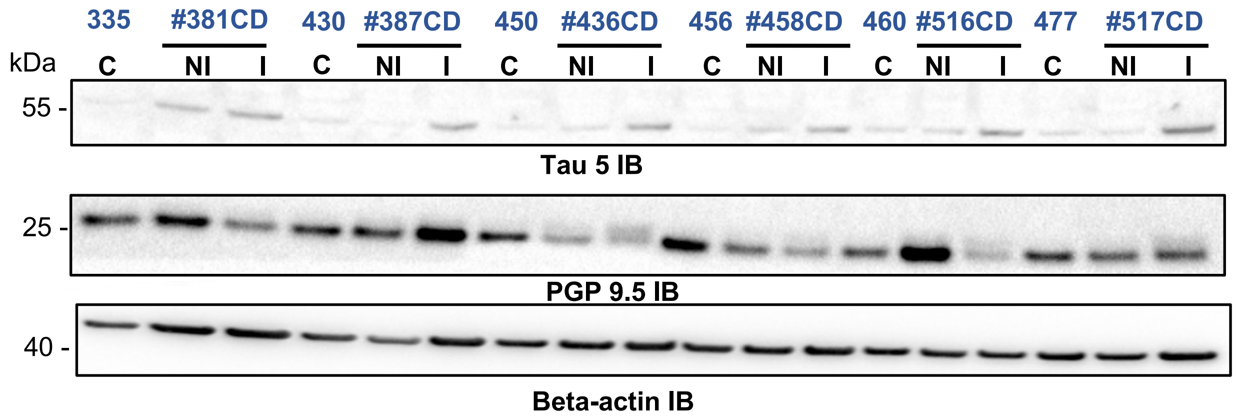


Raw immunoblots for CD and UC patients and quantification for UC patients

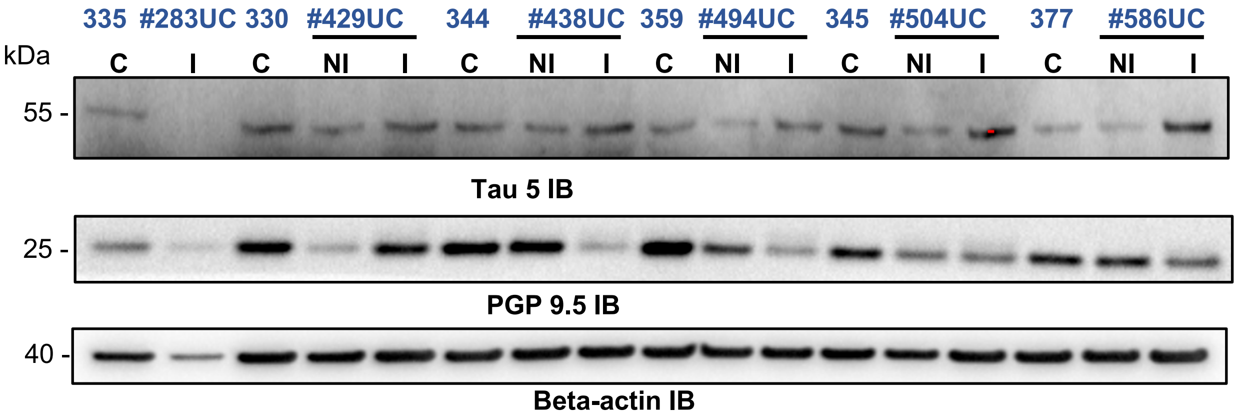


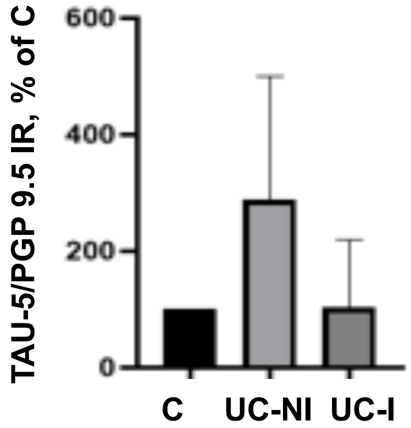

**Supplementary file 3**

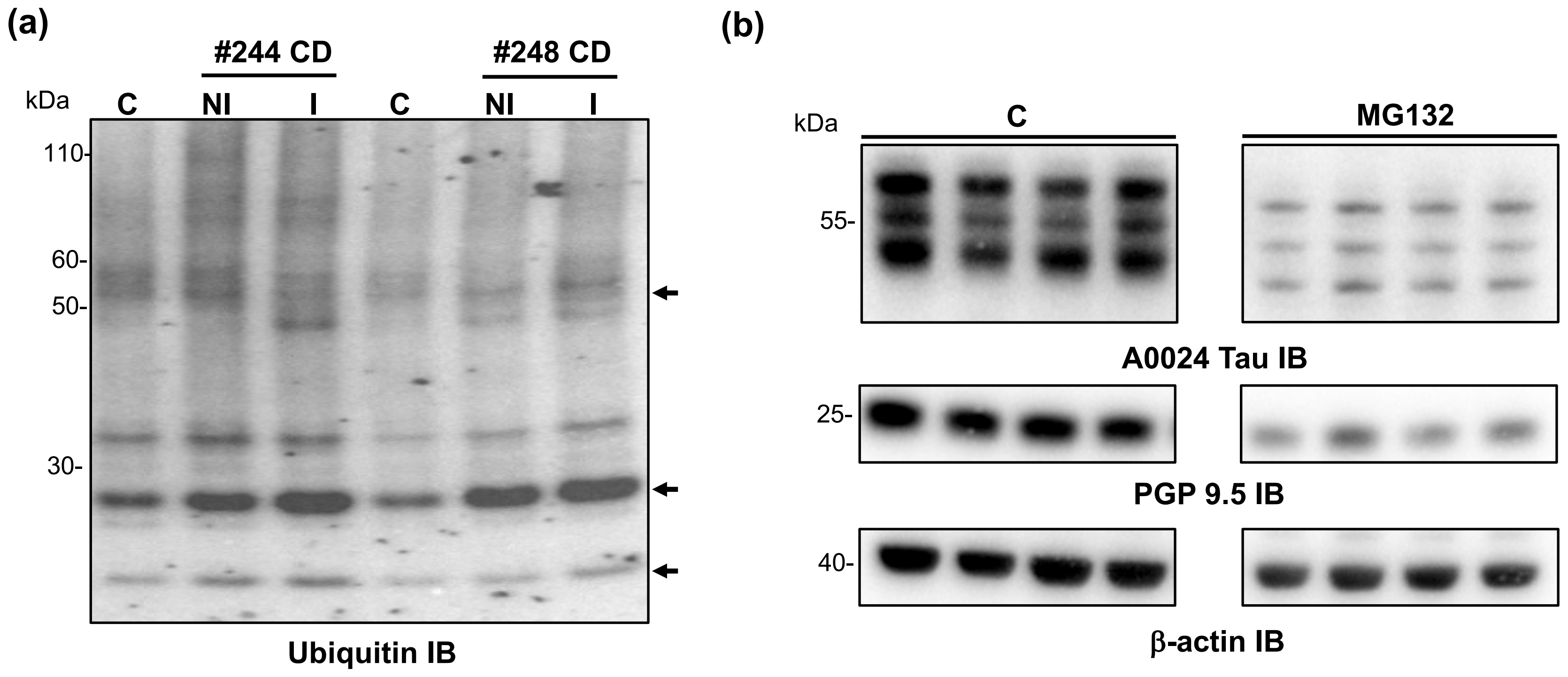


(a) Representative immunoblots of ubiquitin levels in biopsy lysates from CD patients and control subjects. The arrows show some of the bands whose levels varied between CD and controls. (b) Primary culture of rat ENS were treated or not with 10 μM MG132 for 12h and 15 μg of cell lysates were subjected to immunoblot analysis using A0024 Tau, PGP 9.5 (PGP 9.5 IB) and β-actin (β-actin IB) antibodies.
